# Clonucopya: An end-to-end workflow bridging tumor clonal architecture and drug prioritization

**DOI:** 10.64898/2026.10.08.757690

**Authors:** Guillermo Sánchez-Cid, Carlos León-Ramos, Elena Piñeiro-Yáñez, Gonzalo Gómez-López, Fátima Al-Shahrour

## Abstract

**Motivation:** Intra-tumor heterogeneity limits the effectiveness of anticancer treatments, as distinct clonal populations may respond differently to therapy. Although subclonal inference from sequencing data is well established, translating clonal architecture into actionable drug recommendations remains a major bottleneck.

**Results:** We present Clonucopya, an end-to-end Snakemake workflow that integrates subclonal inference (PyClone-VI, PhyClone) with drug prioritization (PanDrugs2) to link tumor clonal architecture directly to candidate drugs. From genome sequencing data, Clonucopya reconstructs tumor phylogenies, quantifies clone-level prevalence across samples, and ranks FDA/EMA-approved and experimental therapeutic candidates against the somatic mutations detected in each clone. Outputs include clonality plots, per-clone variant and drug tables, and a summary ranking drugs by clonal coverage to highlight broadly active therapeutic candidates. We assess Clonucopya’s utility through a real-world longitudinal, multi-region use case, revealing clinically actionable clonal dynamics across tumor progression.

**Availability:** Clonucopya is available as open-source software with full documentation and tutorials at: https://github.com/cnio-bu/clonucopya

## 1 INTRODUCTION

Intra-tumoral heterogeneity (ITH) arises from the accumulation of genetic and non-genetic alterations during tumor evolution, giving rise to subclonal populations with distinct molecular profiles and drug sensitivities. Under selective pressure from treatment, resistant subclones may expand, driving therapeutic failure and relapse (Dijkstra et al., 2025; Hessey et al., 2026). This dynamic clonal evolution represents a major challenge for precision oncology, as treatment decisions based solely on the dominant tumor clone may overlook therapeutically relevant subclonal populations (Bath et al., 2024; Pérez-González, Bévant and Blanpain, 2023). Genome sequencing has been incorporated into routine cancer diagnosis and management (van Putten *et al*., 2026; Sosinsky *et al*., 2024), driving the development of computational methods for detecting genomic variants (Yi *et al*., 2023) and inferring tumor clonal evolution (Wang *et al*., 2024). These methodologies provide researchers and clinicians with deeper insights into tumor biology and more accurate patient assessment (Schaff *et al*., 2024; Lahtinen *et al*., 2023). Clonal inference methods such as PhyloWGS and PyClone-VI use bulk DNA sequencing data to characterize the clonal architecture of tumor cell populations (Deshwar *et al*., 2015; Gillis and Roth, 2020), but their translation into actionable treatment decisions remains limited.

*In silico* drug prioritization tools systematically match genomic alterations with therapeutic options using curated drug-gene interactions and clinical evidence. Among these, PanDrugs2 integrates multi-source genomic and pharmacological evidence to rank treatment recommendations (Jiménez-Santos *et al*., 2022, 2023). Despite their translational utility, these tools generally rely on a bulk tumor consensus profile rather than on the clonal composition of the tumor, overlooking clone-specific therapeutic vulnerabilities. This limitation is particularly relevant because ‘clonetherapy’, defined as a therapeutic strategy designed to address individualized ITH by targeting both clonal and subclonal alterations (Piñeiro-Yañez *et al*., 2019) and thereby minimize the selective pressure that drives treatment resistance. However, no current computational tool integrates clonal inference with clone-specific drug prioritization.

To address this gap, we present Clonucopya, an automated workflow that integrates subclonal inference with *in silico* drug prioritization to translate tumor clonal architecture into actionable therapeutic recommendations. Clonucopya software is intended for bioinformaticians but generates comprehensive reports, including tables and figures, describing the clonal composition of each sample alongside prioritized drug candidates designed for interpretation by translational researchers. By identifying drug candidates capable of targeting multiple tumor clones, Clonucopya supports the design of therapeutic strategies aimed at minimizing the risk of clonal resistance and relapse.

## 2 FEATURES AND METHODS

Clonucopya has been developed to bridge the translational gap between tumor clonal inference and genomics-driven drug prioritization (Figure 1). The software is executed through a structured sequential workflow, requiring as input a VCF file for single nucleotide variants (SNVs) and a CSV file for copy number variants (CNVs), together with a configuration file (YAML). This file allows an easy set up of parameters and scalability to run multiple analyses in parallel. Additionally, Clonucopya accepts complementary sample information (i.e. sex and tumor purity) provided in a TSV-formatted sample sheet. Optionally, the software can use BAM files to estimate variant allele frequencies (VAFs) and also accepts PyClone-VI formatted input files, allowing users to bypass upstream processing steps thus accommodating datasets with precomputed clonal inference results.

**Figure 1.**
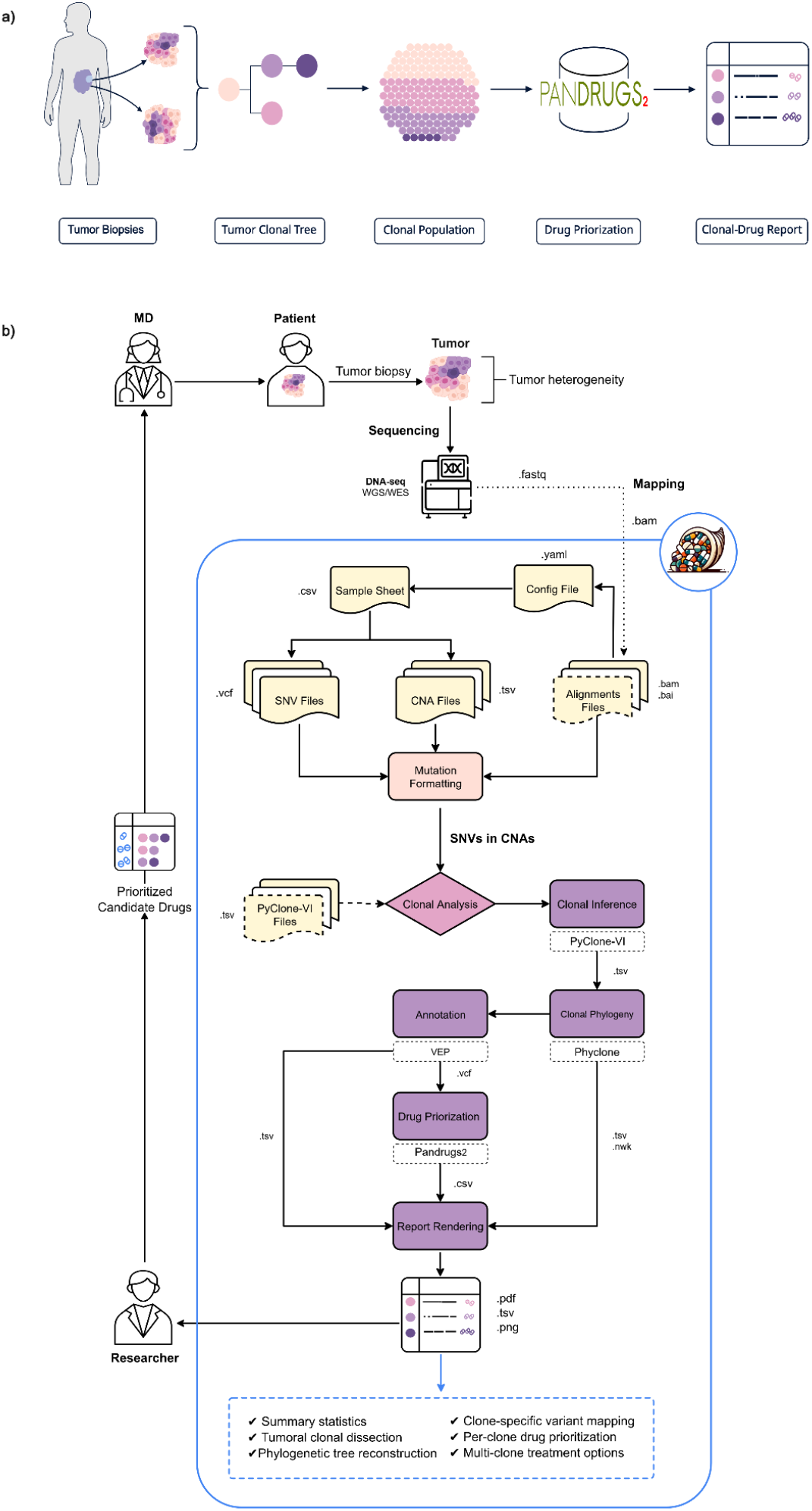
(a) Overview of Clonucopya and (b) schematic representation of the Clonucopya workflow.

Clonucopya supports multi-sample studies enabling both longitudinal and multi-region designs, where multiple paired VCF/CNV inputs from the same study are jointly analyzed to perform pairwise comparisons and reconstruct a unified clonal architecture. Clonucopya intersects SNVs with CNVs and formats the overlapping variants as a PyClone-VI input file, performing clonal inference and clustering the identified clonal populations (Gillis and Roth, 2020). Then, uses PhyClone to compute a phylogenetic tree from these clusters, assigning each cluster to a defined clone within the tumor hierarchy (Roth *et al*., 2025). Somatic variants are functionally annotated using Ensembl-VEP (McLaren *et al*., 2016). The annotated, clone-resolved variants are subsequently queried against PanDrugs2 to retrieve a prioritized ranking of candidate drugs for each clone (Jiménez-Santos *et al*., 2023). The main output is a self-contained PDF report summarizing the clonal and therapeutic landscape. For each sample, it displays per-sample quality metrics, clonal phylogenetic tree, clone distribution across samples and clone-resolved VAF heatmaps. Clonucopya’s tabular outputs include: (i) variant-per-clone table detailing the somatic alterations assigned to each clonal population; (ii) drug-per-clone table listing the therapeutic options prioritized for each individual clone considering PanDrugs GScore and DScore (Piñeiro-Yañez *et al*., 2018); and (iii) a drug summary table across clone groups, providing a comprehensive therapeutic overview of the tumor. Clonucopya also provides additional features to enhance flexibility, robustness, and reproducibility. The pipeline incorporates quality control steps to assess the integrity of input files, including optional BAM files. Furthermore, reproducibility can be improved by specifying a fixed random seed for PyClone-VI analyses. Users may also optionally include indels prior to clonal inference.

### 2.1. Implementation details

Clonucopya is a Python (v3.12.8) command-line workflow for Linux/UNIX and macOS implemented in Snakemake v9.20.0 (Mölder *et al*., 2021). Its modular design supports both end-to-end and stepwise execution, enabling scalable, reproducible, and parallel analyses on HPC environments. Software dependencies are managed through Conda v24.11.3. Clonal inference and phylogenetic reconstruction are performed using PyClone-VI v0.1.6 and PhyClone v0.7.0, respectively. Variants are annotated with Ensembl VEP v116.1 using GRCh38 human genome annotation release 116 as the default reference. Drug candidate prioritization is performed using the PanDrugs2 methodology, which relies on PanDrugsDB, a database integrating 23 primary data sources and comprising more than 74,000 drug-gene associations, involving 4,642 genes and 14,659 unique compounds. The drug prioritization reports include results generated by default with PanDrugs2 “clinical mode” settings, which prioritize drug candidates based on their clinical evidence and therapeutic relevance (Jiménez-Santos *et al*., 2023).The workflow validates input files, supports independent sample processing and dynamic parallelization, and generates detailed logs. Failed tasks preserve intermediate files and are automatically retried or resubmitted when required. X

#### 2.1.1. Benchmarking

Clonucopya has been executed in a 9 compute node cluster with 728 CPU cores (6 × 52 core/512Gb RAM and 3 × 64core/768Gb RAM). Using a simulated dataset of 10 samples from the ICGC-TCGA DREAM Somatic Mutation Calling-Tumor Heterogeneity (SMC-Het) Challenge (Salcedo *et al*., 2020) with paired SNV and CNV calls but no BAM files, the workflow generates the final report in 12h 50min using 8 cores. We further tested performance using PyClone-VI input files from the same dataset, with sample mutations counts ranging from 103 to 105. Running *pvi-start* execution mode with default parameters, Clonucopya completed the analysis in 13 h 4min using 8 cores.

## 3 USE CASES

To demonstrate both the technical validity and practical utility of Clonucopya, two datasets were analyzed: i) the simulated P10 tumor from the SMC-Het Challenge, used to validate Clonucopya implementation and the clonal inference workflow, and ii) five tumor samples from a lung adenocarcinoma patient (CRUKP7127) enrolled in the PEACE cohort (Hessey *et al*., 2026), used to illustrate the identification and prioritization of clone-specific therapeutic strategies in a real patient with multi-region, longitudinal sampling. Complete step-by-step analysis code for both use cases along with the corresponding PDF reports, are available in the Supplementary Files and in the Clonucopya tutorials (https://github.com/cnio-bu/clonucopya/tree/main/tutorial).

### 3.1. DREAM Challenge P10 synthetic tumor

We tested Clonucopya’s technical implementation using P10, a synthetic tumor sample available as supplementary material for PyClone-VI and via Synapse (project ID: syn2813581). The input files contain a total of 8,080 SNVs located in regions affected by CNVs. The PyClone-VI implementation within the Clonucopya workflow detected all 7,708 mutations reported in the original PyClone-VI analysis of P10, supporting the reliability of our implementation (Suppl Figure 1). The remaining 372 SNVs had a median VAF=0.089, consistent with subclonal variants or technical noise (VAF <0.1). The reconstructed clonal architecture reveals a linear phylogenetic tree comprising six distinct clones (c0-c5; clonal prevalences: c0=12%, c1=6%, c2=25%, c3=24%, c4=14%, and c5=19%). Clone c3 showed the highest mutational burden (3,411 mutations), followed by c0 (2,359), c1 (1,158), c5 (1,118), c4 (31), and c2 (3). Drug prioritization identified 39 candidate drugs targeting at least three clones (Suppl File 1). Among multi-clone candidates, fostamatinib, a SYK inhibitor approved for chronic immune thrombocytopenia, emerged as a promising repurposing candidate in cancer (Hu *et al*., 2025). Standard chemotherapies were also predicted to target multiple clones, with docetaxel covering c0/c3/c5, doxorubicin, cyclophosphamide, and fluorouracil covering c0/c1/c3, and cisplatin covering c0/c3.

### 3.2. Longitudinal multi-region analysis of PEACE patient CRUKP7127

PEACE (Posthumous Evaluation of Advanced Cancer Environment) is a multi-centre study characterizing tumor evolution through multi-region and longitudinal sampling of advanced cancers (Hessey *et al*., 2026). We applied Clonucopya to 5 multi-region and longitudinal tumor samples from a 71-year-old female PEACE patient with lung adenocarcinoma (ID: CRUKP7127). Four samples were obtained from the primary tumor (SU_T1-R1, SU_T1-R2, SU_T1-R3, SU_T1-R4) and one sample from a pathologist-confirmed locoregional relapse involving lymph node (BR_LN01). The patient underwent surgical resection with adjuvant therapy (cisplatin+vinorelbine) and following nodal relapse received carboplatin+atezolizumab+pemetrexed. Clonucopya identified substantial ITH, with ten clones (c0–c9) derived from ancestral clone c0. Clone c0 branched into c1, c2 and c3, with c3 giving rise to two divergent lineages: c4–c7 and c8–c9 (Suppl File 2). Clonal composition differed markedly between samples, with clone c7 dominating SU_T1-R2 (97%) and clone c5 predominating in SU_T1-R1 (41%), consistent with site-specific clonal selection during tumor evolution. Drug prioritization identified a missense mutation in PIK3CA (3:179221146:G>A; VAF 0.029-0.055) recently reported as an oncogenic variant in ClinVar (ID:VCV000376476.64), present in c2 and shared across clones 2, 5, 6, 7 and 9, as the most therapeutically actionable alteration (GScore = 0.799). PanDrugs2 prioritized the PI3K inhibitor alpelisib as the top therapeutic candidate (DScore = 0.920), based on direct target and pathway-member interactions. Importantly, c3, shared across all five samples, carried a frameshift mutation in UFL1, a gene previously implicated in chromosome instability and tumorigenesis, for which no targeted therapies are currently available (Li *et al*., 2026). For truncal clone c0, present in all five samples, Clonucopya prioritized marimastat, the CDK inhibitors PHA-793887 and ronciclib, and 5-fluorouracil, reflecting a potential broad actionability of this early clonal event. Clone c1, detected exclusively in SU_T1-R3 and carrying a BAZ2B missense variant, was prioritized for doxorubicin and tamoxifen (Dscores = 0.822 and 0.812, respectively), both supported by biomarker-level evidence (Suppl Fig).

The patient’s clinical course provides a context for the clonal landscape inferred by Clonucopya. Adjuvant cisplatin was predicted to be active against clones c2, c6 and c9, whereas the truncal clone c0 and its descendant c3 were not associated with cisplatin sensitivity; both clones could have persisted in the subsequent lymph node relapse sample (BR_LN01: c0, c3, c5, c6), together with c6. Although causality cannot be established from this single case, the observation is compatible with recurrence from a c0→ c3→ c5/c6 lineage not directly targeted by adjuvant therapy. None of the administered drugs (cisplatin, vinorelbine, atezolizumab or pemetrexed) targeted PIK3CA, which PanDrugs2 prioritized as the top therapeutic candidate (alpelisib, DScore = 0.920). This discrepancy between the empirical treatment received and the genomically-informed prioritization illustrates the potential utility of Clonucopya integrating clonal evolution analysis into treatment selection to guide therapeutic strategies.

## 4 CONCLUSIONS

We developed Clonucopya, a Snakemake workflow integrating tumor clonal dissection, evolution analysis, and drug prioritization to target ITH. Using bulk DNA sequencing data, Clonucopya reconstructs tumor phylogenies across related samples and identifies clone-specific drug vulnerabilities. The workflow is designed for bioinformaticians and produces intuitive reports for translational and clinical researchers. Key applications include prioritizing treatments based on clone-specific genomic alterations across single-sample, multi-region, and longitudinal data, as well as the identification of resistant clones and evolutionary trajectories associated with therapeutic failure. Clonucopya insights can support evolution-informed precision oncology, in which treatment selection accounts for the clonal composition and trajectory of a tumor rather than relying solely on its dominant or most recently sampled genomic profile. Clonucopya source code, full documentation and step-by-step tutorials are accessible under a GPL-3.0 license at https://github.com/cnio-bu/clonucopya

## ACKNOWLEDGEMENTS

The authors thank Bioinformatics Unit staff, E. Arroba and M. Robledo from the Endocrine Hereditary Cancer group at CNIO for beta testing and fruitful discussions.

## Author contributions

G.S-C developed Clonucopya and wrote the manuscript. C.L-R and E.P-Y collaborated in the implementation of the code. G.G-L supervised the study, contributed to conceptualization, methodology, formal analysis, and wrote the manuscript. F.A-S. supervised the study, secured funding, managed the project, and contributed to manuscript review and editing. All authors reviewed and approved the final manuscript.

## Data availability

Clonucopya source code, documentation, and links to all datasets used in this study are publicly available at https://github.com/cnio-bu/clonucopya, including the PyClone-VI supporting datasets hosted on Zenodo (https://zenodo.org/records/4268826).

---

**Supplementary Figure 1.**
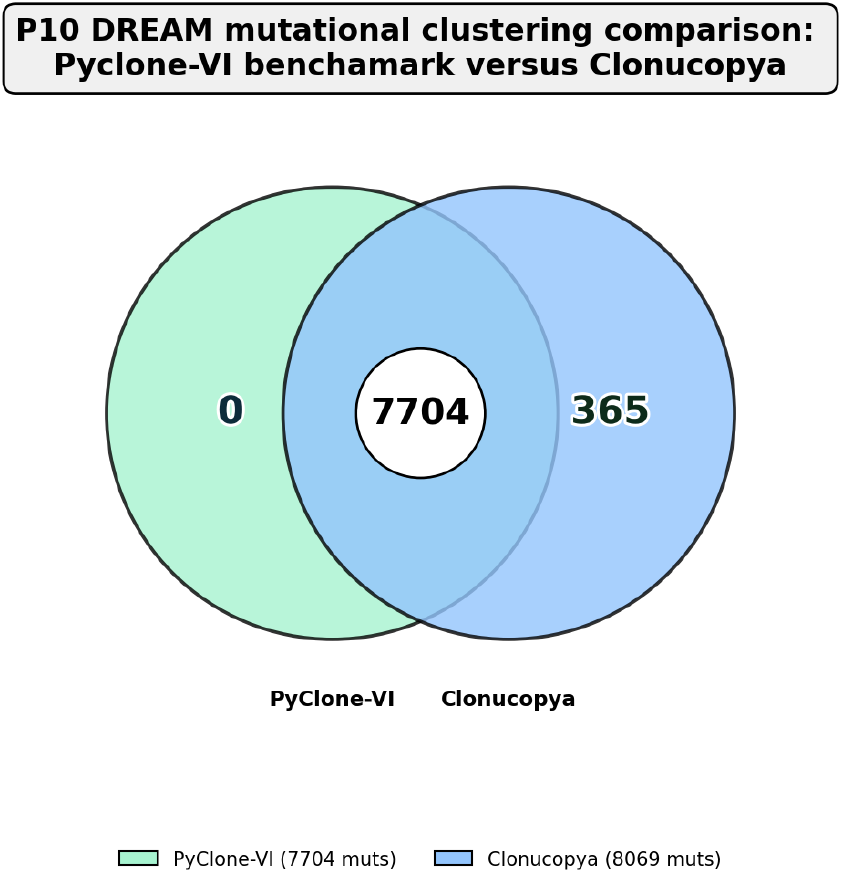
Comparison of mutations contained in sample P10 from Dream Challenge processed dataset and Clonucopya dataset obtained from raw data (simulated SNV and CNV variant calling).

**Supplementary Figure 2.**
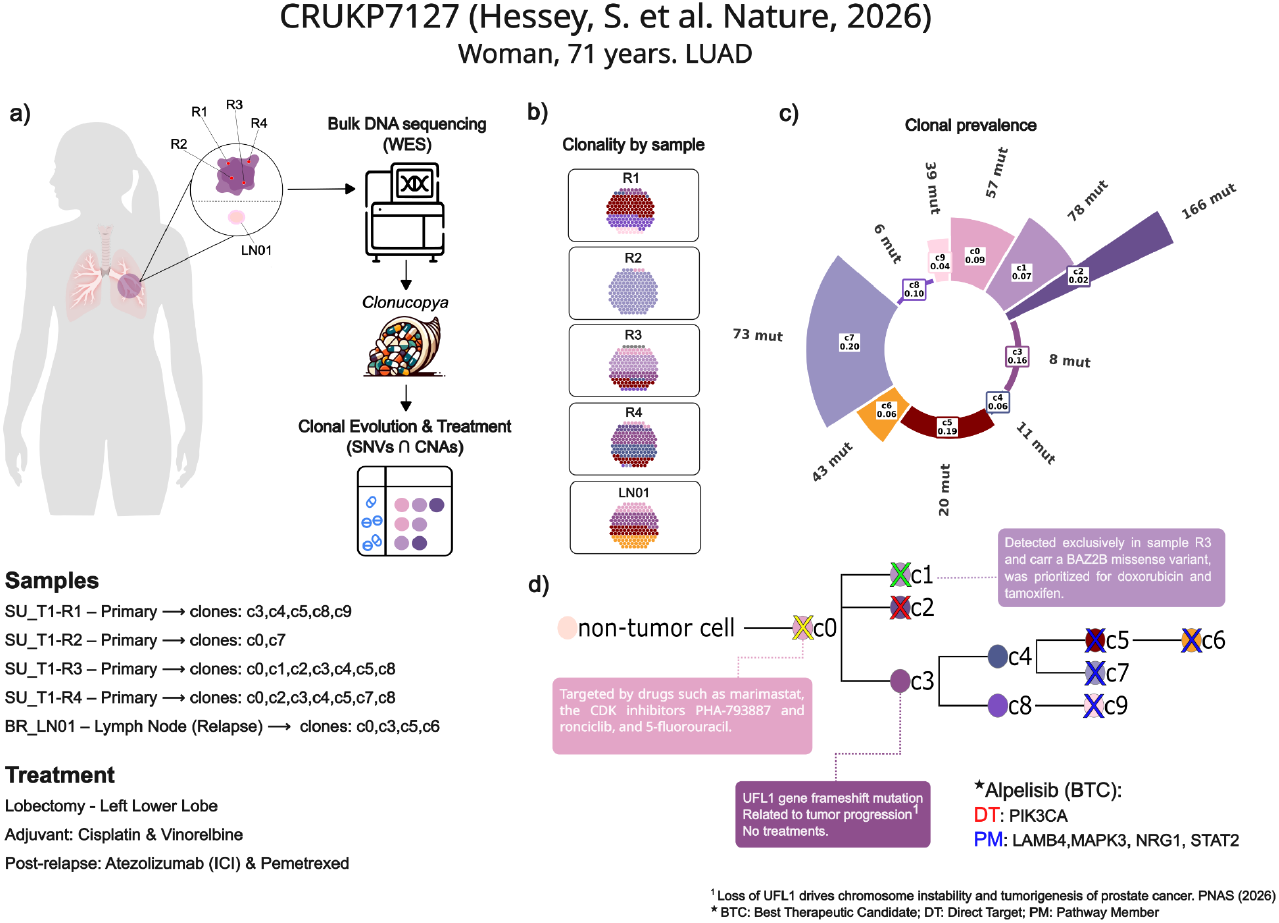
Clonal architecture and evolutionary history of metastatic lung adenocarcinoma in patient CRUKP7127 from the PEACE cohort. (a) Overview of the CRUKP7127 case, a 71-year-old woman with lung adenocarcinoma (LUAD), including multi-region sampling of the primary tumour (R1–R4) and longitudinal sampling of a lymph-node relapse (LN01), together with the patient’s treatment history (lobectomy followed by adjuvant cisplatin/vinorelbine and, after relapse, atezolizumab plus pemetrexed). Whole-exome sequencing (WES) data were subsequently analysed with Clonucopya to reconstruct clonal evolution and identify clone-specific therapeutic opportunities from SNVs and CNAs. (b) Clonal composition inferred for each tumour sample. (c) Clonal prevalence by sample, showing the relative abundance of inferred clones and their associated mutational burden. (d) Reconstructed phylogenetic relationships between tumour clones, showing the emergence and diversification of clones c0–c9 from a common non-tumour ancestral population. Potential therapeutic treatments are suggested by PanDrugs2 based on the genomic alterations identified in each clone, with selected drug–target associations indicated in the phylogenetic tree.

